# Systemic inflammation is a minor contributor to remnant cholesterol atherogenicity

**DOI:** 10.1101/2024.07.18.604203

**Authors:** Jordan M. Kraaijenhof, Marije J. Kerkvliet, Nick S. Nurmohamed, Aldo Grefhorst, Jeffrey Kroon, Nicholas J. Wareham, G. Kees Hovingh, Erik S.G. Stroes, S. Matthijs Boekholdt, Laurens F. Reeskamp

## Abstract

**Background:** Both plasma levels of remnant cholesterol and low-density lipoprotein cholesterol (LDL-C) levels are independent risk factors for atherosclerotic cardiovascular disease. However, only remnant cholesterol has consistently been associated with systemic inflammation. The extent to which inflammation mediates the effect of remnant cholesterol on major adverse cardiovascular events (MACE) remains unclear.

**Methods and Results:** This study included 16,445 participants without prior atherosclerotic cardiovascular disease from the EPIC-Norfolk cohort, with a mean age of 58.8±9.1 years, of which 9,357 (56.9%) were women. Every 1 mmol/L higher remnant cholesterol was associated with 29.5% higher hsCRP levels (95% Confidence Interval (CI): 22.1, 37.4, p<0.001), whereas LDL-C was not significantly associated with hsCRP levels in the fully adjusted model. Additionally, each 1 mmol/L higher remnant cholesterol was associated with a hazard ratio (HR) of 1.31 (95% CI: 1.14, 1.50, p<0.001) for MACE, compared to a HR of 1.21 (95% CI: 1.13, 1.31, p<0.001) for LDL-C. Mediation analysis showed that hsCRP mediated 5.9% (95% CI: 1.2, 10.6%, p<0.001) of the effect of remnant cholesterol on MACE, whereas hsCRP did not mediate the effect of LDL-C.

**Conclusions:** Plasma remnant cholesterol levels are independently associated with systemic inflammation and cardiovascular events. Inflammation, as measured with hsCRP, contributed minorly to the association between remnant cholesterol and MACE. This underscores the need to address both remnant cholesterol and systemic inflammation separately in the clinical management of cardiovascular disease.

Graphical abstract:
The study assessed the relationship between remnant cholesterol, systemic inflammation, and MACE risk in 16,445 participants free from atherosclerotic cardiovascular disease from the EPIC-Norfolk cohort. Every 1 mmol/L higher remnant cholesterol was associated with 29.5% higher hsCRP levels (95% CI: 22.1, 37.4, p<0.001), while LDL cholesterol was not significantly associated with hsCRP levels. Additionally, each 1 mmol/L higher remnant cholesterol was associated with a HR of 1.31 (95% CI: 1.14, 1.50, p<0.001) for MACE, compared to a HR of 1.21 (95% CI: 1.13, 1.31, p<0.001) for LDL-C. hsCRP mediated 5.9% (95% CI: 1.2, 10.6%, p<0.001) of the effect of remnant cholesterol on MACE, while it did not mediate the effect of LDL cholesterol. LDL: low-density lipoprotein cholesterol, HR: hazard ratio, CI: confidence interval, MACE: major adverse cardiovascular events.

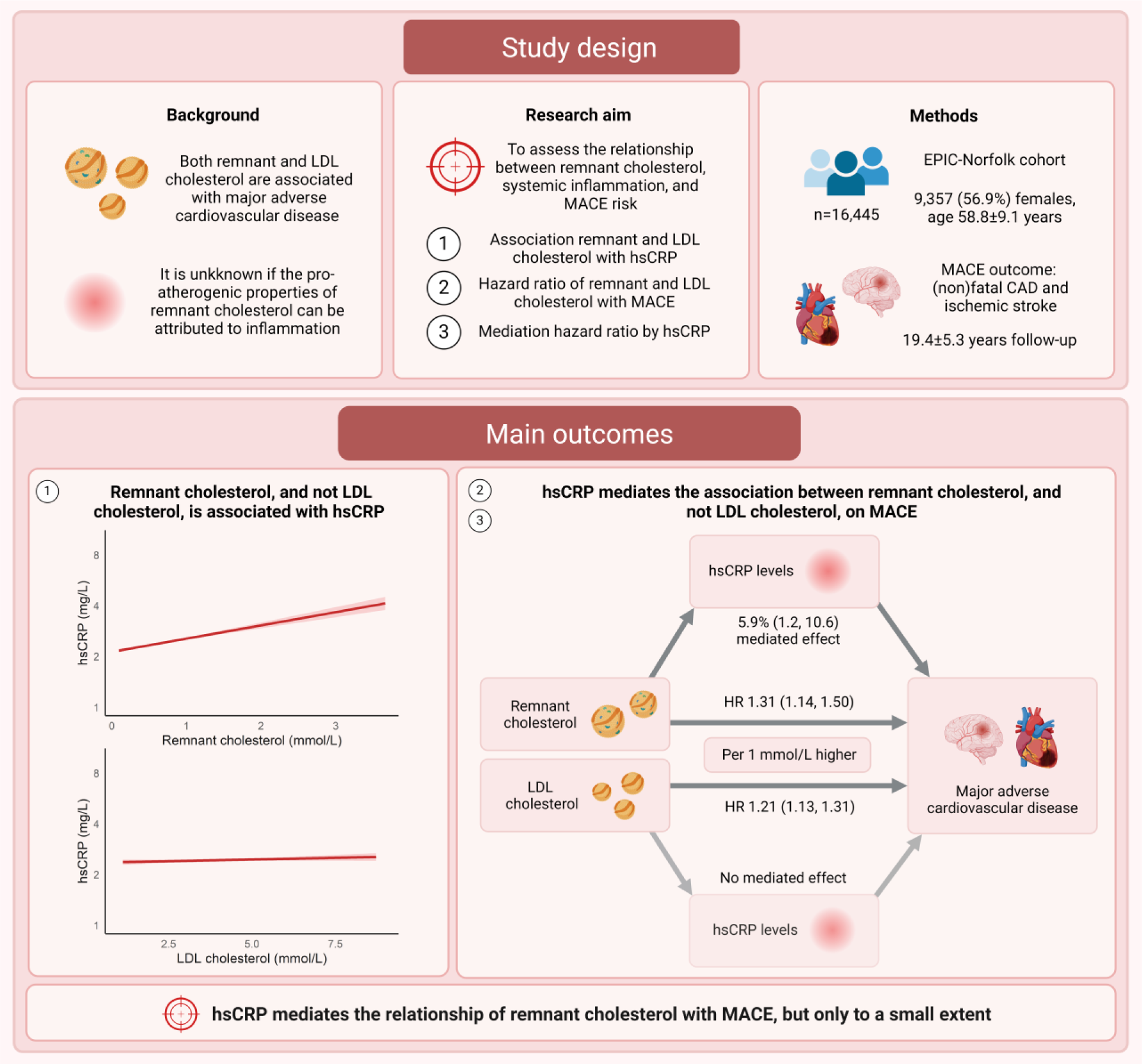

## Introduction

Apolipoprotein B (apoB)-containing particles are causal partakers during initiation and progression of atherogenesis.(1,2) Triglyceride-rich apoB-containing particles (TRLs) are produced in the liver as very-low-density-lipoproteins (VLDL) and are subsequently metabolized to VLDL-remnant particles, part of which are further processed to low-density lipoproteins (LDL) through lipoprotein-lipase mediated lipolysis.(3) Several metabolic disorders such as obesity and insulin resistance increase VLDL production and impair lipolysis, resulting in accumulation of remnant particles in the circulation.(4) The cholesterol content of remnant particles, commonly referred to as remnant cholesterol, as well as remnant particle related proteins such as apolipoprotein C-III (apoC-III), have been independently associated with major adverse cardiovascular disease (MACE).(5–8).

While large observational and genetic studies unanimously underscore a predominant role for the concentration of apoB-containing particles in causing MACE (9–13), recent genetic studies have implied that remnant cholesterol may have an even bigger impact on cardiovascular risk compared with cholesterol in LDL particles (LDL cholesterol), suggesting a specific pro-atherogenic effect beyond apoB-particle concentration.(14) It was suggested that this observation could be attributed to the impact of remnant particles on inflammation.(15) Experimental studies have underscored the inflammatory capacity of the large free-fatty-acid load in remnant particles inducing inflammation via lipoprotein lipase (LPL)-induced lipolysis as well as the potential for receptor-independent uptake of TRLs by subendothelial macrophages.(15) Clinical studies corroborated a pro-inflammatory effect of remnant particles, showing a correlation between remnant particles and arterial wall inflammation assessed by FDG-PET/CT as well as systemic inflammation measured by high-sensitivity C-reactive protein (hsCRP) levels.(16–22) It remains to be established what extent of remnant cholesterol’s atherogenic risk is mediated by this pro-inflammatory effect.

In this study, we aimed to assess the contribution of systemic inflammation assessed as hsCRP to the MACE risk of remnant and LDL cholesterol in the large prospective EPIC-Norfolk cohort. We also evaluated if this effect is confounded by the presence of metabolic syndrome factors, known to be independently associated with systemic inflammation. Lastly, we detailed this relationship further by investigating particle characteristics (i.e. particle size and apoC-III levels) of remnant cholesterol containing particles and their effect on inflammation.

## Methods

### Study population

The study included participants from the European Prospective Investigation into Cancer and Nutrition (EPIC)-Norfolk prospective cohort study free of atherosclerotic cardiovascular disease at time of in enrollment, involving 25,639 men and women between the ages of 40 and 79, all from general practices in Norfolk, United Kingdom.(23) Participants were recruited between 1993 and 1998 and were followed for over 25 years with check-up visits and questionnaires, with follow-up data available up to 31^st^ of March 2016. A nested case-control study with available plasma samples within the EPIC-Norfolk cohort was designed in 2004 to study the association between plasma biomarkers and CVD. Cases were study participants who did not have prevalent CVD at baseline, but did develop CAD during follow-up through the end of 2003.(24) Controls, matched on sex, age (±5 years), and enrollment date (±3 months) in a 2:1 ratio, were selected from participants who remained free of cardiovascular disease during follow-up. In this sub-study, VLDL particle size and apoC-III levels measurements were performed (details provided below). Ethical approval was granted by the Norwich District Health Authority Ethics Committee, with all participants having provided written informed consent.

### Cardiovascular disease and other disease definitions

In the current study, the endpoint was a composite of MACE, comprised of (non)fatal coronary artery disease (CAD) (ICD-10; I20–I25) and (non)fatal ischemic stroke (ICD-10; I63 and I65–I66), as done previously in EPIC Norfolk (24) Metabolic syndrome was defined as the presence of 3 out of 5 characteristics: increased waist circumference (≥102 cm for men and ≥88 cm for women), elevated triglyceride levels (≥1.7 mmol/L), high systolic blood pressure (≥130 mmHg) or high diastolic blood pressure (≥85 mmHg), hyperglycemia defined as elevated HbA1c of 42 mmol/mol (6%) and above, and low HDL-C levels (<1.03 mmol/L for men and <1.30 mmol/L for women).(28) DM was defined as either use of diabetic mediation or an HbA1c of 48 mmol/mol (6.5%) and above.

### Laboratory measurements

Baseline non-fasting blood samples were analyzed for total cholesterol (TC), high-density lipoprotein cholesterol (HDL-C), triglycerides and apolipoprotein B (apoB) using a RA 1000 auto-analyzer (Bayer Diagnostics, Basingstoke, United Kingdom). LDL-C levels were estimated with the NIH Sampson equation (25) in order to achieve more precision in estimating LDL-C levels at elevated triglyceride levels and enhanced accuracy in calculating VLDL-C levels compared to the Friedewald formula.(26) Remnant cholesterol was estimated by subtracting HDL-C and LDL-C from total cholesterol levels.(27) hsCRP levels were measured using an Olympus AU640 Chemistry Immuno Analyzer (Olympus Diagnostics, Watford, United Kingdom). Participants with hsCRP levels above 10 mg/L were excluded from the analysis to eliminate individuals with acute infectious or systemic inflammatory diseases.

In the nested case-control study, the size and particle concentrations of VLDL, comprising of small, medium, and large particles, were measured using an automated 400-megahertz proton nuclear magnetic resonance (NMR) spectroscopic assay.(29) VLDL particle size was quantified in nmol/L and categorized based on the diameter: small particles 27-35 nm, medium particles 35-60 nm, and large particles >60 nm.(29) Plasma concentrations of apoC-III were measured using a chemiluminescent enzyme-linked immunoassay (Abgent, San Diego, United States of America).(30)

### Study outcomes

The primary outcome was the hsCRP mediated association between remnant or LDL cholesterol with future MACE. Secondary outcomes included the association between concentration of small, medium and large VLDL particles and apoC-III levels with hsCRP levels, investigated within the nested case-control sub-study.

### Statistical analysis

Normally distributed data and non-normally distributed data are presented as mean ± standard deviation (SD) and as median ± interquartile range [IQR], respectively. Categorical data is reported as absolute numbers and percentages. Demographic, clinical and biochemical characteristics are provided for the complete cohort and stratified by remnant cholesterol quartiles. An ANOVA test was performed for normally distributed continuous variables, while a Kruskal–Wallis test was used for non-normally distributed variables. Correlations were examined with the Pearson’s rank correlation test for non-normally distributed data.

To explore the associations between remnant cholesterol, LDL-C, and log_2_ transformed hsCRP levels, linear regression analysis was used with additional adjustment for age and sex. Next, we corrected the association for the presence of metabolic syndrome factors diabetes mellitus (DM), body mass index (BMI) and systolic blood pressure (SBP). Next, we corrected these models for current smoking. In a final model, we corrected for LDL-C in the remnant cholesterol model and vice versa.

Next, Cox proportional hazards models were used to calculate hazard ratios (HRs) per 1 mmol/L or 1 standard deviation increment in remnant and LDL cholesterol in relation to MACE.(31) The initial model was adjusted for age, sex, diabetes mellitus, BMI, systolic blood pressure, and current smoking. An additional model was then constructed with further adjustment for apoB levels and LDL-C in the remnant cholesterol model and vice versa.

Mediation analysis was conducted to investigate the role of hsCRP in mediating the relationship between remnant or LDL cholesterol levels and cardiovascular disease events using the CMAverse package (32). This method utilizes Cox proportional hazard modelling with nonparametric bootstrapping with 1,000 runs to estimate the total, direct, and indirect effects. Estimates and confidence intervals (CIs) below 0% were truncated to 0%. Sensitivity analyses were performed in subgroups stratified for sex (men and women) and metabolic syndrome (0 to 2 MS features versus 3 or more).

To ascertain the independent effects of remnant cholesterol or LDL-C on cardiovascular events and their mediation by hsCRP levels, we conducted a discordance analysis using Cox proportional hazards modeling with four groups based on the median values of remnant (RC) and LDL-C levels: low RC/low LDL-C as the reference group (remnant and LDL cholesterol < median cohort levels), low RC/high LDL-C, high RC/low LDL-C and high RC/high LDL-C. These analyses were fully adjusted for age, sex, BMI, DM, SBP, and smoking status.

In the nested case-control cohort study, we examined the association between remnant cholesterol, remnant particle-related measures, including concentrations of small, medium, and large VLDL particles and apoC-III levels, and log2-transformed hsCRP levels. We used a linear regression model, adjusting for age, sex, DM, BMI, SBP, smoking status, and LDL-C. To validate the role of systemic inflammation in these remnant-related measures, we applied logistic regression models to assess the odds ratio for incident CAD. These models were adjusted for age, sex, DM, BMI, SBP, smoking status, and an additional model including LDL-C.

Statistical significance was set at a p-value of less than 0.05 and all statistical analyses were performed using RStudio version 4.3.2. (R Foundation, Vienna, Austria).

## Results

In the EPIC-Norfolk cohort study, lipid and hsCRP levels were available in 18,445 individuals. After excluding patients with atherosclerotic cardiovascular disease at baseline and participants with hsCRP levels greater than 10 mg/L, the final analysis included a total of 16,445 individuals (Table 1), with a mean age of 58.8±9.1 years, of whom 9,357 (56.9%) were women. The median levels of remnant and LDL cholesterol were 0.61 [0.42, 0.88] mmol/L and 4.0 (1.0) mmol/L, respectively.

**Table 1.** Baseline characteristics.

| Characteristics | Overall cohort | Q1 remnant<br>cholesterol<br><0.42 mmol/L | Q2 remnant<br>cholesterol 0.42-0.61 mmol/L | Q3 remnant<br>cholesterol 0.61-0.88 mmol/L | Q4 remnant<br>cholesterol<br>>0.88 mmol/L | p-value |
| --- | --- | --- | --- | --- | --- | --- |
| Number of patients | 16,445 | 4,112 | 4,113 | 4,110 | 4,110 |  |
| Age (years) | 58.8 (9.1) | 56.0 (8.9) | 58.7 (9.2) | 60.0 (9.1) | 60.5 (8.7) | **<0.001** |
| Male sex | 9,357 (56.9%) | 2,937 (71.4%) | 2,492 (60.6%) | 2,110 (51.3%) | 1,818 (44.2%) | **<0.001** |
| Current smoker | 1,805 (11.1%) | 410 (10.0%) | 439 (10.8%) | 457 (11.2%) | 499 (12.2%) | **0.013** |
| Diabetes mellitus | 288 (1.8%) | 53 (1.3%) | 54 (1.3%) | 77 (1.9%) | 104 (2.5%) | **<0.001** |
| Lipid-lowering medication | 174 (1.1%) | 28 (0.7%) | 38 (0.9%) | 55 (1.3%) | 53 (1.3%) | **0.009** |
| Antihypertensive medication | 2,581 (15.7%) | 408 (9.9%) | 573 (13.9%) | 716 (17.4%) | 884 (21.5%) | **<0.001** |
| BMI (kg/m<sup>2</sup>) | 25.6 [23.5, 28.0] | 24.1 [22.3, 26.3] | 25.1 [23.1, 27.3] | 26.1 [24.1, 28.5] | 27.1 [25.1, 29.5] | **<0.001** |
| Waist circumference (cm) | 87.1 (12.1) | 80.6 (10.5) | 85.3 (11.3) | 89.4 (11.4) | 93.5 (11.1) | **<0.001** |
| Systolic blood pressure (mmHg) | 134.6 (18.2) | 129.0 (17.6) | 133.2 (18.0) | 136.5 (17.8) | 139.7 (17.7) | **<0.001** |
| Diastolic blood pressure (mmHg) | 82.0 (11.1) | 78.8 (10.7) | 81.2 (10.9) | 83.0 (11.0) | 85.2 (10.9) | **<0.001** |
| Apolipoprotein B (g/L) | 1.0 (0.2) | 0.9 (0.2) | 0.9 (0.2) | 1.0 (0.2) | 1.1 (0.3) | **<0.001** |
| Total cholesterol (mmol/L) | 6.2 (1.1) | 5.7 (1.0) | 5.9 (1.0) | 6.3 (1.1) | 6.8 (1.1) | **<0.001** |
| LDL cholesterol (mmol/L) | 4.0 (1.0) | 3.7 (1.0) | 3.9 (1.0) | 4.2 (1.0) | 4.4 (1.0) | **<0.001** |
| HDL cholesterol (mmol/L) | 1.4 (0.4) | 1.7 (0.4) | 1.5 (0.4) | 1.3 (0.4) | 1.2 (0.3) | **<0.001** |
| Triglycerides (mmol/L) | 1.5 [1.1, 2.1] | 0.8 [0.7, 0.9] | 1.3 [1.1, 1.4] | 1.8 [1.6, 1.9] | 2.6 [2.3, 3.1] | **<0.001** |
| VLDL-cholesterol (mmol/L) | 0.7 [0.5, 1.0] | 0.4 [0.3, 0.4] | 0.5 [0.5, 0.6] | 0.8 [0.7, 0.9] | 1.2 [1.1, 1.5] | **<0.001** |
| Remnant cholesterol (mmol/L) | 0.6 [0.4, 0.9] | 0.3 [0.3, 0.4] | 0.5 [0.5, 0.6] | 0.7 [0.7, 0.8] | 1.1 [1.0, 1.4] | **<0.001** |
| High-sensitivity C-reactive protein (mg/L) | 1.4 [0.7, 2.8] | 0.9 [0.5, 1.9] | 1.3 [0.7, 2.6] | 1.6 [0.8, 3.1] | 1.9 [1.0, 3.5] | **<0.001** |

Baseline characteristics for the complete cohort and per remnant cholesterol quartile. Normally distributed variables are reported as mean (SD), non-normally distributed variables as median ± interquartile range [IQR], and categorical variables as number (%). These were analyzed using a one-way ANOVA, a Kruskal–Wallis test, or a Chi<sup>2</sup> test, respectively. BMI: body mass index, LDL: low-density lipoprotein, HDL: high-density lipoprotein, VLDL: very-low-density lipoprotein.

**Table 2.** MACE hazard ratios for remnant and LDL cholesterol.

|  | **Hazard ratio (95% CI)**<br>Per 1 mmol remnant<br>cholesterol | **Hazard ratio (95% CI)**<br>Per 1 mmol LDL<br>cholesterol |
| --- | --- | --- |
| Model 1 – Age and sex | 1.70 (1.56, 1.84) | 1.19 (1.16, 1.23) |
| Model 2 – Model 1 + MS components | 1.38 (1.22, 1.57) | 1.19 (1.13, 1.23) |
| Model 3 – Model 2 + current smoking | 1.36 (1.20, 1.55) | 1.19 (1.13, 1.26) |
| Model 4 – Model 3 + apoB, LDL-C/RC | 1.31 (1.14, 1.50) | 1.21 (1.13, 1.31) |

Cox proportional modeling was used to determine hazard ratios for remnant and LDL cholesterol on future MACE. MACE was defined as (non)fatal coronary artery disease and (non)fatal ischemic stroke. Model 1 was age and sex adjusted, model 2 was additionally adjusted for MS factors BMI, DM and SBP, model 3 additionally for current smoking and model 4 for apoB and LDL-C in model A and remnant cholesterol in model B. LDL: low-density lipoprotein, MACE: major adverse cardiovascular disease, BMI: body mass index, DM: diabetes mellitus, SBP: systolic blood pressure, apoB: apolipoprotein B.

The prevalence of obesity, increased waist circumference and hypertension were higher in those with higher remnant cholesterol levels (Table 1). Moreover, compared to those in the lowest remnant cholesterol quartile, study participants in the highest quartile had higher LDL-C and lower HDL-C levels (4.4±1.0 vs. 3.7±1.0 mmol/L, p<0.001, and 1.2±0.3 vs. 1.7±0.4 mmol/L, p<0.001, respectively). The baseline characteristics according to the metabolic syndrome scoring are provided in Table S1.

### The association between remnant cholesterol, LDL cholesterol and plasma hsCRP levels

Every 1 mmol/L higher remnant cholesterol was associated with 72.7% (95% CI: 66.0, 80.0, p<0.001) higher hsCRP levels after adjustment for age and sex. This association was attenuated after adjustment for MS components, with 1 mmol/L higher remnant cholesterol levels corresponding to 33.1% (95% CI: 25.6, 41.1, p<0.001) higher hsCRP levels. After further adjustment for current smoking, hsCRP levels were 30.4% (95% CI: 23.0, 38.2, p<0.001) higher per 1 mmol/L increase in remnant cholesterol. Lastly, after LDL-C adjustment, a 1 mmol/L higher remnant cholesterol corresponded to 29.5% (95% CI: 22.1, 37.4, p<0.001) higher hsCRP levels (Table S2 and Figure 1A).

**Figure 1.**
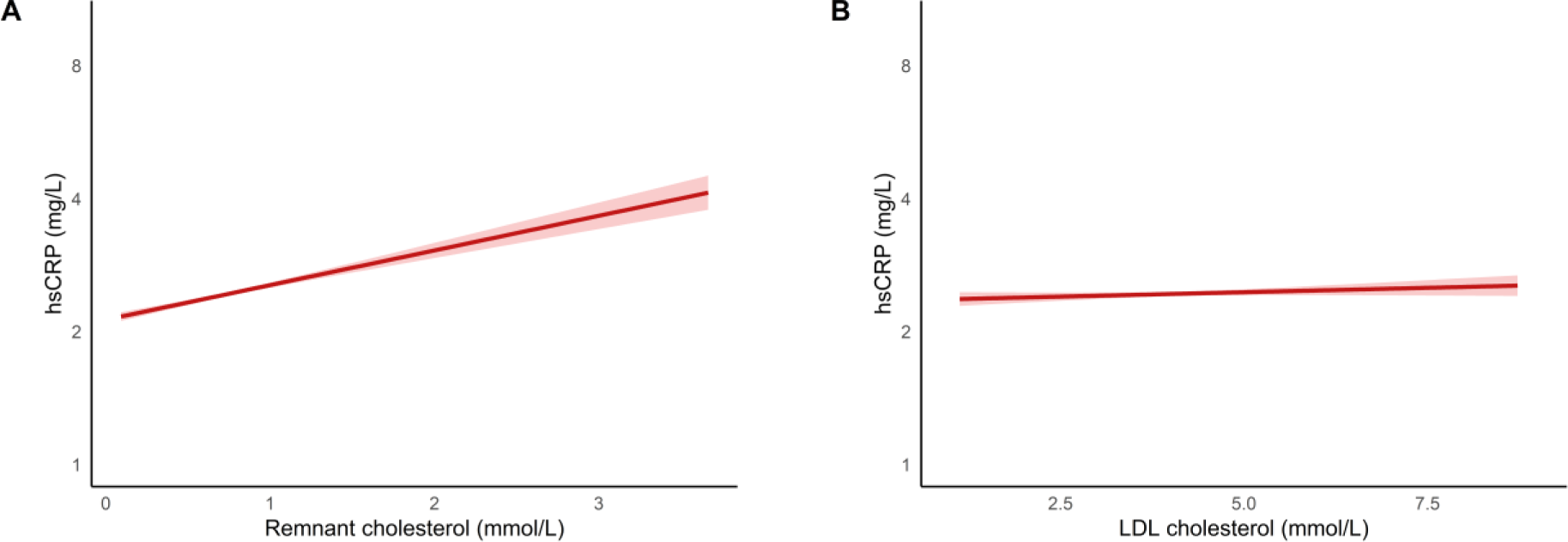
Relationship between remnant cholesterol, LDL cholesterol and hsCRP levels. Visualization of linear regression analyses for A) remnant cholesterol and B) LDL-C plasma levels on hsCRP levels (on a log2 scale) in a multivariable adjusted model, corrected for age, sex, DM, BMI, SBP, current smoking and LDL-C for (A) and remnant cholesterol for (B). Light red areas indicate 95% confidence interval of the fitted regression line. LDL: low-density lipoprotein, DM: diabetes mellitus, BMI: body mass index, SBP: systolic blood pressure.

LDL cholesterol showed a markedly weaker association with hsCRP levels. For every 1 mmol/L increment in LDL-C levels, hsCRP levels were 7.5% (95% CI: 6.0, 9.1, p<0.001) higher in the age and sex-adjusted model, 3.8% (95% CI: 1.5, 6.1, p<0.001) higher in the MS component adjusted model and 3.3% (95% CI: 1.1, 5.6, p=0.004) higher in the model additionally adjusted for smoking. After adjustment for remnant cholesterol, no significantly higher hsCRP per 1 mmol/L higher LDL-C was observed. (Table S2 and Figure 1B). Results per one SD higher remnant and LDL cholesterol are provided in Table S2.

### The impact of remnant and LDL cholesterol on the risk for cardiovascular disease

During a median follow-up period of 19.4±5.3 years, 3,466 MACE events occurred. One mmol/L higher remnant cholesterol was associated with a HR of 1.70 (95% CI: 1.56, 1.84, p<0.001) for MACE in the age and sex adjusted model. In the fully adjusted model following adjustment for MS components, current smoking, LDL-C and apoB, the hazard ratio was 1.31 (95% CI: 1.14, 1.50, p<0.001) per 1 mmol/L higher remnant cholesterol (Figure 2). Mediation analysis showed that 5.9% (95% CI: 1.2, 10.6%, p<0.001) of the relationship between remnant cholesterol levels and MACE was mediated by hsCRP (Figure 3). One mmol/L higher LDL cholesterol was associated with a hazard ratio of 1.19 (95% CI: 1.16, 1.23, p<0.001) for MACE in the age and sex adjusted model. In the fully adjusted model, one mmol/L higher LDL-C resulted in a hazard ratio of 1.21 (1.13, 1.31, p<0.001) (Figure 2). In contrast to remnant cholesterol, no significant mediation by hsCRP was observed (Figure 3). Because the metabolic syndrome is associated with systemic inflammation, we investigated if these results were different for those with or without 3 or more metabolic syndrome features. For the 1.434 participants with MS, the hazard ratio was not significant, at 1.03 (95% CI: 0.82, 1.31, p=0.776), with no significant mediation (Table S3). Conversely, the 5,506 individuals with no MS, the hazard ratio for MACE was 1.40 (95% CI: 1.13, 1.75, p<0.001) per 1 mmol/L higher remnant cholesterol of which 11.0% (95% CI: 1.3-23.4, p<0.001) was mediated by hsCRP.

**Figure 2.**
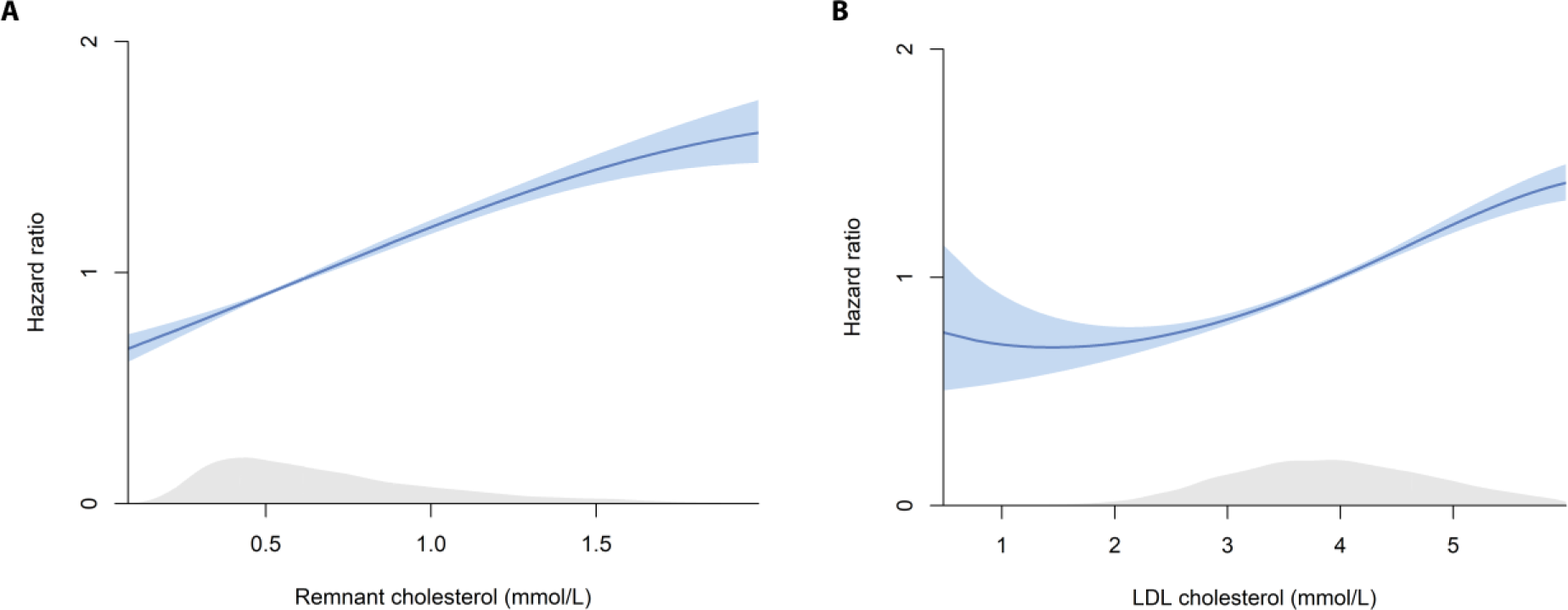
Association of remnant and LDL cholesterol with MACE. Cox proportional hazard ratios for remnant (A) and LDL cholesterol (B) on future MACE. MACE was defined as (non)fatal coronary artery disease and (non)fatal ischemic stroke. Results are shown for the multivariable adjusted model, corrected for age, sex, DM, BMI, SBP, current smoking, apoB and LDL-C in A and remnant cholesterol in B. Grey area indicates the density of measurements of remnant (A) and LDL cholesterol (B). LDL: low-density lipoprotein, MACE: major adverse cardiovascular disease, DM: diabetes mellitus, BMI: body mass index, SBP: systolic blood pressure, apoB: apolipoprotein B.

**Figure 3.**
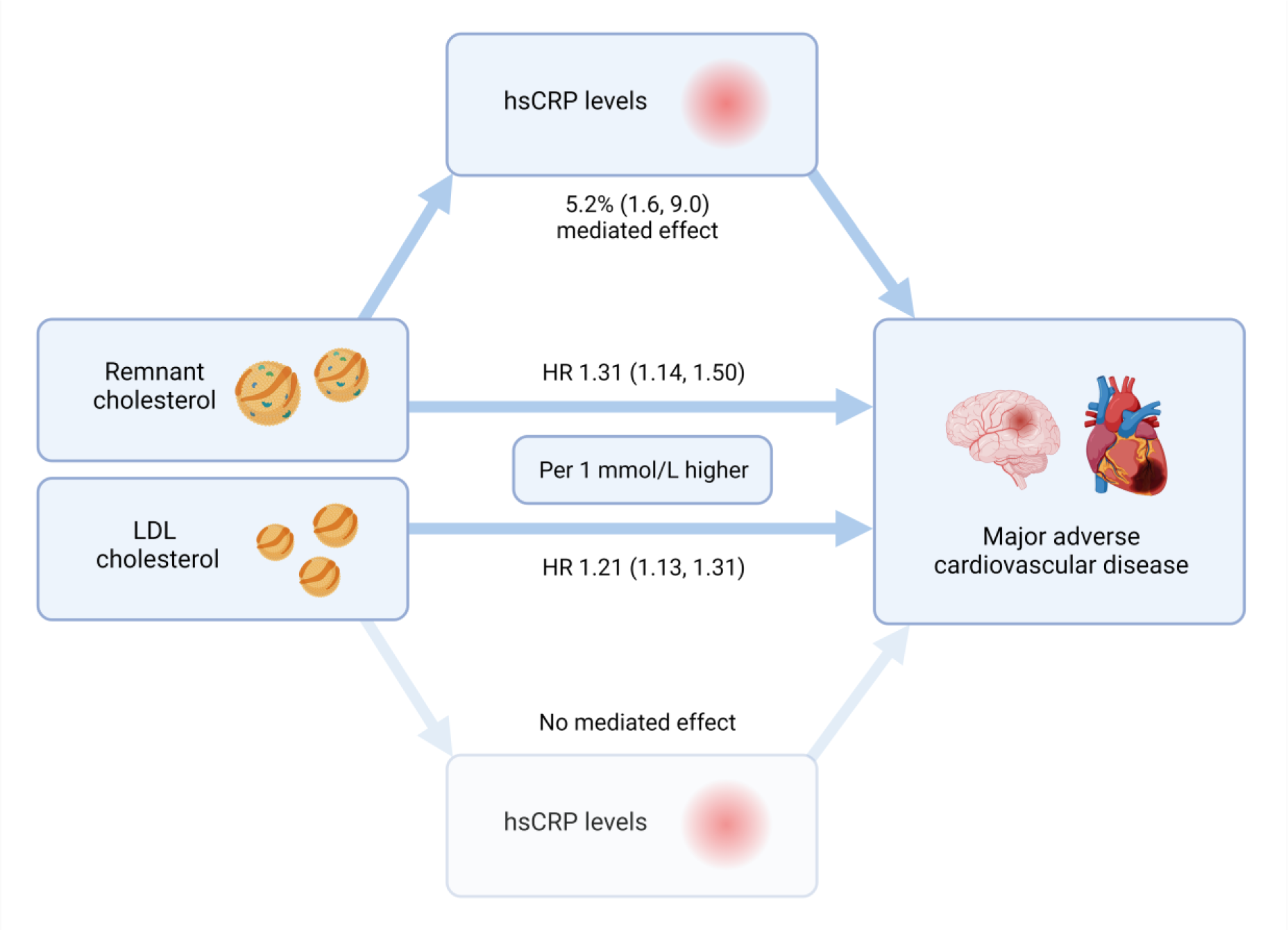
Mediation analysis of hsCRP levels of remnant and LDL cholesterol on MACE. Mediation analysis between the proportion effect of hsCRP in the association between remnant and LDL cholesterol with major adverse cardiovascular disease. hsCRP: high-sensitivity C-reactive protein, LDL: low-density lipoprotein, HR: hazard ratio. Created with Biorender.com.

Lastly, we found no sex difference in the magnitude of the effect of hsCRP on the association between remnant cholesterol and MACE. In men, the hazard ratio of 1 mmol/L higher remnant cholesterol for MACE was 1.29 (95% CI: 1.09, 1.53, p=0.004; Table S3), with the mediated effect of hsCRP being 8.8% (95% CI: 3.4, 18.2, p<0.001). For women, the hazard ratio was 1.28 (95% CI: 1.02, 1.60, p=0.031), and the mediated effect was non-significant.

### Analysis of discordance: for the impact of remnant and LDL cholesterol on MACE and the mediation by hsCRP levels

Four groups, based on the median of remnant and LDL cholesterol levels, were constructed to investigate discordancy in MACE associated risk. The low RC/low LDL-C served as reference group (remnant cholesterol and LDL cholesterol below median cohort levels).

Compared to the reference group, the HR for MACE was 1.22 (95% CI: 1.02, 1.45, p=0.031, Table S4) in the low RC/high LDL-C group and 1.20 (95% CI: 1.01, 1.42, p=0.036) in the high RC/low LDL-C group (Figure 4 and Table S4) in the fully adjusted model. For the high RC/high LDL-C group, the HR was 1.46 (95% CI: 1.26, 1.70, p<0.001). Mediation by hsCRP was observed in the high RC/low LDL-C group with 2.8% (1.5, 4.1, p<0.001) and in the high RC/high LDL-C group with 2.0% (1.0, 3.0, p<0.001) (Figure 4 and Table S4). No mediation was observed in the low RC/high LDL-C group.

**Figure 4.**
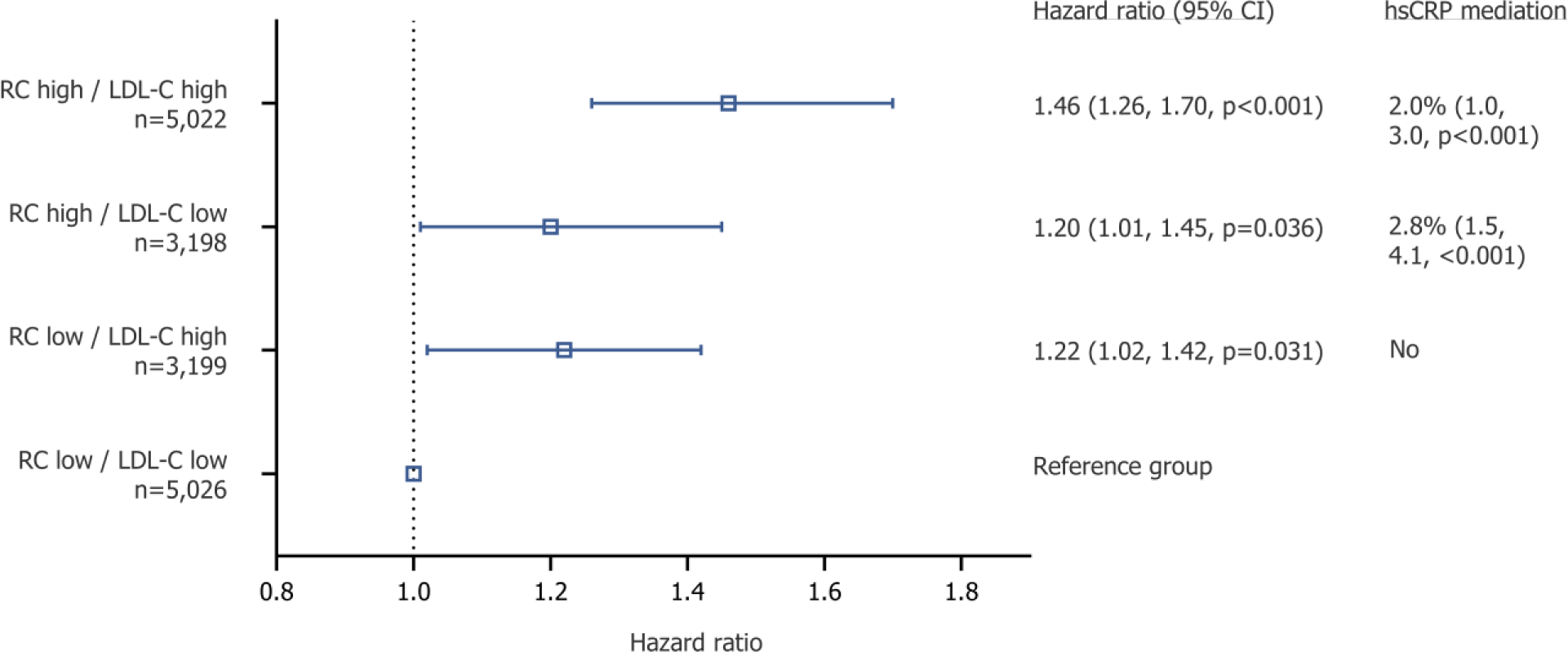
Hazard ratio and mediation analysis in discordant analysis. Discordance analyses based on the median values of remnant cholesterol (0.6 mmol/L) and LDL cholesterol (4.0 mmol/L) levels; the low RC/low LDL-C (reference group), high RC/low LDL-C, low RC/high LDL-C, and high RC/high LDL-C group. Cox proportional hazard ratios calculated for future MACE, defined as (non)fatal coronary artery disease and (non)fatal ischemic stroke and its mediation by hsCRP levels. Results are shown for the multivariable adjusted model, corrected for age, sex, DM, BMI, SBP and current smoking. LDL: low-density lipoprotein, MACE: major adverse cardiovascular disease, hsCRP: high-sensitivity C-reactive protein, DM: diabetes mellitus, BMI: body mass index, SBP: systolic blood pressure.

### The impact of remnant cholesterol and particle properties on hsCRP in the nested case-control study

To further detail the effect of remnant particles characteristics on the relationship between remnant cholesterol and hsCRP, we performed a sub study in the nested case-control EPIC-Norfolk study for which hsCRP levels were available in 2,987 participants which comprised of 961 cases and 2,063 controls. A comparison of baseline characteristics between the cohort study and case-control subset is presented in Table S5. Remnant cholesterol levels showed the following correlation with triglycerides (Pearson correlation coefficient (ρ) = 0.99, p<0.001), small VLDL particles (ρ = 0.20, p<0.001), medium VLDL particles (ρ = 0.58, p<0.001, large VLDL particles (ρ = 0.85, p<0.001) and with apoC-III (ρ = 0.34, p<0.001; Figure S1).

In accordance with the longitudinal cohort analysis, remnant cholesterol was associated with systemic inflammation. One SD higher remnant cholesterol was associated with 11.4% (95% CI: 7.4, 15.5, p<0.001) higher hsCRP levels in a model adjusted for age, sex, DM, BMI, SBP, current smoking, and LDL cholesterol (Table 3). One standard deviation higher concentrations of medium and large VLDL particle were significantly associated with 4.9% (95% CI: 1.0, 9.0, p=0.014) and 13.9% (95% CI: 9.5, 18.4, p<0.001) higher hsCRP levels respectively. Small VLDL size concentrations were not associated with hsCRP levels. One standard deviation higher plasma apoC-III levels was associated with 8.9% (95% CI: 4.7, 13.3, p<0.001) higher hsCRP levels.

**Table 3.** Relationship between remnant cholesterol (related markers) and hsCRP.

|  | **Higher hsCRP (95% CI)**<br>Per 1 mmol increase | **p-value** |
| --- | --- | --- |
| Remnant cholesterol | 11.4% (7.4, 15.5) | **<0.001** |
| Small VLDL size | -0.1% (-4.6, 4.6) | 0.95 |
| Medium VLDL size | 4.9% (1.0, 9.0) | **0.014** |
| Large VLDL size | 13.9% (9.5, 18.4) | **<0.001** |
| Apolipoprotein C-III | 8.9% (4.7, 13.3) | **<0.001** |

Linear regression analysis for remnant cholesterol and standardized higher small, medium and large VLDL concentration and apoCIII levels on hsCRP levels. All models are multivariable adjusted for age, sex, DM, BMI SBP, smoking status and LDL-C. hsCRP: high-sensitivity C-reactive protein, VLDL: very-low-density lipoprotein, DM: diabetes mellitus, BMI: body mass index, SBP: systolic blood pressure, LDL: low-density lipoprotein.

One SD higher remnant cholesterol was associated with an odds ratio of 1.19 (1.09, 1.30, p<0.001) for CAD, adjusted for age, sex, DM, BMI, SBP, current smoking, and LDL-C with 7.6% (3.0, 15.0), p<0.001) mediation by hsCRP levels (Table S6). Each standard deviation higher medium and large VLDL particle concentration was associated with an odds ratio of 1.11 (1.01, 1.21, p=0.002) and 1.08 (0.98, 1.19, p=0.104) for CAD respectively, with no significant mediation by hsCRP (Table S6). Each standard deviation increase in apoC-III levels was associated with an odds ratio of 1.13 (95% CI: 1.03, 1.25, p=0.012) for CAD, with no significant mediation by hsCRP.

## Discussion

In the present study, we confirm that plasma levels of remnant cholesterol are associated with hsCRP levels, whereas hsCRP only modestly mediated the association between remnant cholesterol and cardiovascular events (Figure 5). The association between remnant cholesterol and hsCRP is markedly attenuated by metabolic syndrome features and could potentially be attributed to large, triglyceride loaded, particles carrying the apolipoprotein apoC-III. In contrast to remnant cholesterol, LDL-C was only weakly correlated with hsCRP, and hsCRP did not significantly mediate LDL-associated cardiovascular disease risk.

The results of our study align with previous studies showing that remnant cholesterol is associated with cardiovascular disease beyond apoB and LDL cholesterol. In a study by Quispe and coworkers, remnant cholesterol levels were associated with the risk of cardiovascular events in 17,532 primary prevention individuals, independent of traditional cardiovascular risk factors, LDL-C, and apoB.(33) This is corroborated by a pooled cohort study of intravascular ultrasound trials in patients with known coronary artery disease, which showed that on-treatment remnant cholesterol levels were associated with coronary atheroma progression after adjusting for apoB.(5) Conversely, these results contrast with other studies suggesting that cardiovascular risk associated with remnant cholesterol can be completely attributed to the fact that it is just cholesterol carried in apoB containing lipoproteins that can become trapped in the arterial wall.(9–13)

One explanation for our observation that remnant cholesterol is associated with ASCVD beyond apoB could lie in its inflammatory effects. Multiple large-scale studies indicated that both observational and genetically determined plasma levels of remnant cholesterol are associated with higher hsCRP levels indicative of an increased systemic inflammatory state.(19,21,22) Our study adds to this knowledge by showing a strong correlation between remnant cholesterol levels and hsCRP levels and remnant cholesterol levels and ASCVD in another large cohort. Through our mediation analyses, we aimed to assess which proportion of the observed effect of remnant and LDL cholesterol on cardiovascular event risk could be attributable to hsCRP. Only 5% of the MACE risk associated with remnant cholesterol was found to be mediated through hsCRP. This is consistent with the discordance analyses suggesting that, although there is a strong association between remnant cholesterol and systemic inflammation, the pro-inflammatory effects of remnant cholesterol as quantified by hsCRP are unlikely major drivers of the increased cardiovascular event risk. The weak mediation effect of hsCRP in the association of large VLDL particles and apoC-III with coronary artery disease further substantiates this finding and suggests that the independent association of remnant cholesterol (related markers) with systemic inflammation might be of limited clinical relevance.

This raises the question whether other factors contribute to remnant cholesterol atherogenicity. First, it could be that the pro-inflammatory effects of remnant particles are effectuated on a cellular level and localized to the arterial wall, which is not (fully) reflected by plasma hsCRP levels. Experimental studies have shown that the triglyceride and free fatty acid load in these particles are susceptible to lipolysis, which can exert pro-inflammatory effects on endothelial cells and subendothelial macrophages.(10,37) Clinical studies have found that remnant cholesterol levels are independently associated with vascular inflammation using FDG-PET/CT imaging, which was not correlated with hsCRP levels (20)(38). Changes in atherosclerotic inflammation in the arterial wall independent from hsCRP changes is supported by the observation that LDL-C lowering with PCSK9 inhibition significantly reduces vascular inflammation without affecting hsCRP levels.(39) Second, on a per particle basis, remnant particles contain more cholesterol and have a longer residence time in the vasculature compared with LDL particles.(3) ApoC-III could play a particular role in this process as in vitro studies have shown that ApoC-III on triglycerice-rich lipoproteins enhance the binding to biglycans; negatively charged glycosaminoglycans present in lesions, potentially increasing vascular retention of atherogenic lipoproteins.(45) And third, remnant particles are more effective at inducing macrophage foam cell formation compared with LDL particles, as they can initiate receptor-independent uptake in macrophages without requiring oxidative modification hallmarking LDL particle uptake. Lastly, remnant cholesterol may be a marker of an underlying risk factor for ASCVD. For example, a recent study reported that approximately 80% of the MACE risk associated with remnant cholesterol was in fact mediated by insulin resistance.(46) Although we corrected for metabolic syndrome components, we did not measure insulin resistance which thus could be an underlying causal factor.

### Limitations

Our study has several limitations that warrant further discussion. First, the low number of participants with diabetes and those receiving lipid-lowering treatments restricts the applicability of our findings to more contemporary primary prevention populations.(47,48) Secondly, blood was withdrawn in the non-fasted state, leading to elevation of remnant cholesterol levels which may have impacted our observations and limit the generalizability to the fasting state. Third, remnant cholesterol levels have not been measured directly, but were calculated using the NIH Sampson formula. Yet, it has demonstrated a very good correlation with remnant cholesterol levels measured via ultracentrifugation.(26,49,50) Last, the genetic and ethnic homogeneity of the European cohort may limit the applicability of our findings to other regions and patient groups.

## Conclusions

Remnant cholesterol is independently associated with systemic inflammation and atherosclerotic cardiovascular events, even after correction for apoB. However, inflammation only mediates the association between remnant cholesterol and MACE for a minor part. These findings emphasize the need to address both remnant cholesterol and systemic inflammation separately in the clinical management of cardiovascular disease.

## Acknowledgements

The authors gratefully acknowledge the study participants and staff of the EPIC-Norfolk study.

